# Uncovering Alterations in Cancer Epigenetics via Trans-Dimensional Markov Chain Monte Carlo and Hidden Markov Models*

**DOI:** 10.1101/2023.06.15.545168

**Authors:** Farhad Shokoohi, Saeedeh Hajebi Khaniki

## Abstract

Epigenetic alterations are key drivers in the development and progression of cancer. Identifying differentially methylated cytosines (DMCs) in cancer samples is a crucial step toward understanding these changes. In this paper, we propose a trans-dimensional Markov chain Monte Carlo (TMCMC) approach that uses hidden Markov models (HMMs) with binomial emission, and bisulfite sequencing (BS-Seq) data, called `DMCTHM`, to identify DMCs in cancer epigenetic studies. We introduce the Expander-Collider penalty to tackle under and overestimation in TMCMC-HMMs. We address all known challenges inherent in BS-Seq data by introducing novel approaches for capturing functional patterns and autocorrelation structure of the data, as well as for handling missing values, multiple covariates, multiple comparisons, and family-wise errors. We demonstrate the effectiveness of `DMCTHM` through comprehensive simulation studies. The results show that our proposed method outperforms other competing methods in identifying DMCs. Notably, with `DMCTHM`, we uncovered new DMCs and genes in Colorectal cancer that were significantly enriched in the Tp53 pathway.

## 1 Introduction

Epigenetic alterations, such as DNA methylation, histone modifications, and chromatin remodeling, play a critical role in the development and progression of cancer. Such alterations can lead to changes in gene expression and thereby contribute to oncogenesis, tumor growth, and metastasis. In particular, aberrant DNA methylation, which involves the addition of a methyl group to the cytosine residue in CpG dinucleotides, has been widely implicated in cancer initiation and progression. Therefore, identifying differentially methylated cytosines (DMCs) or regions (DMRs) in cancer samples is an essential step toward understanding the underlying epigenetic changes and developing novel therapeutic strategies.

Colorectal cancer (CRC), as the third most common cancer worldwide and the second cause of cancer-related deaths, is a significant global health concern with several risk factors including age, family health history, inflammatory bowel disease, and lifestyle choices (Sung et al., 2021). Unfortunately, many CRC patients remain asymptomatic until the disease has advanced, posing a significant challenge for early detection. However, recent studies have shown that DNA methylation is crucial in the development and progression of CRC (Ashktorab and Brim, 2014). Profiling differentially methylated genes (DMG) in CRC samples compared to normal tissue could identify potential targets for diagnostic or therapeutic interventions. Moreover, DNA methylation analysis could enhance the accuracy of CRC diagnosis and prognosis beyond current staging methods. For instance, *SEPT9, NDRG4, VIM, APC, SFRP1, SFRP4*, and *SFRP5* are the most important CRC-related DMGs (Mueller and Győrffy, 2022). These findings offer tremendous promise for developing epigenetic-based diagnostic and prognostic tools that could lead to more effective and targeted treatments. Therefore, further exploration of epigenetic modifications, such as DNA methylation, is warranted.

Bisulfite sequencing (BS-Seq) has enabled genome-wide DNA methylation profiling, revolutionizing the study of epigenetic changes. However, analyzing BS-Seq data presents several challenges, including capturing functional methylation patterns and autocorrelation structure, dealing with heavy missing values and low read-depth, accounting for multiple covariates, and controlling for multiple comparisons and family-wise errors. Various statistical methods have been proposed for identifying DMCs/DMRs, ranging from classical linear models to advanced machine learning algorithms. In what follows we give a brief overview of these methods.

### 1.1 A Brief Literature Review

DMC/DMR identification methods utilize various statistical models, algorithms, tests, and family-wise error controls. However, not every method uses all of these components, and their application varies between methods. Supplementary Table S1 provides a list of the methods along with their characteristics. Below, we provide a brief overview.

Several DMC/DMR methods utilize Hidden Markov Models (HMMs) for analyzing methylation data. Among others, Shokoohi et al. (2019) introduced `DMCHMM`, which utilizes HMMs in conjunction with the EM and MCMC algorithms to smooth methylation data for individual subjects, followed by a general mixed model to account for covariates and detect DMCs while controlling the family-wise error. `ComMet` (Saito and Mituyama, 2015), Bisulfighter (Saito et al., 2014), `HMMDM` (Yu and Sun, 2016), `HMMFisher` (Sun and Yu, 2016) and `HMMDMR` (Ji, 2019) employ 3-State HMMs, while Molaro et al. (2011) utilize a 2-State HMM to identify differential methylation. Chen et al. (2021) (`BSDMR`) and Shen et al. (2017) (`DMRMark`) apply non-homogeneous HMMs. Other HMM-based methods include `MethPipe` (Song et al., 2013) and R.L. et al. (2013).

A few methods are based on regression models. For instance, `bumphunter` (Jaffe et al., 2012) uses linear mixed models, while methylKit (Akalin et al., 2012) uses logistic regression. `BSmooth (bsseq)` (Hansen et al., 2012), `RADMeth` (Dolzhenko and Smith, 2014), and `GetisDMR` (Wen et al., 2016) use regression-based models such as local linear regression and beta-binomial regression. The `dmrseq` (Korthauer et al., 2018) method uses a generalized least squares regression model with a nested autoregressive correlated error structure, and `MACAU` (Lea et al., 2015) employs a binomial mixed model.

Bayesian models are used by many methods. Among others, `DSS` (Feng et al., 2014) and `DMCFB` (Shokoohi et al., 2021) apply Bayesian functional regression models. `WFMM` (Lee and Morris, 2015) introduces a Bayesian wavelet-based functional linear mixed model. DMRfinder (Gaspar and Hart, 2017) utilizes Bayesian beta-binomial hierarchical modeling. Some methods focus on distribution, likelihood, or kernel density estimation. For example, MethylSig (Park et al., 2014) uses beta-binomial distribution along with a likelihood ratio test. BiSeq (Hebestreit et al., 2013) applies weighted local likelihood to smooth the data and then applies multiple testing to detect DMRs. ICDMR (Hsiao et al., 2014) applies clustering on logit-transformed beta values using normal mixture models. M3D (Mayo et al., 2014) proposes a kernel-based test for spatially correlated changes in methylation profiles.

Several methods rely solely on various statistical tests, including metilene (Jühling et al., 2016), swDMR (Wang et al., 2015), Defiant (Condon et al., 2018), DMRFusion (Yassi et al., 2018), and SMAP (Gao et al., 2015).

Additionally, machine learning algorithms such as CluBCpG (Scott et al., 2020) and HOME (Srivastava et al., 2019) have recently been employed in this context.

For more information, refer to comprehensive reviews by Laird (2010), Robinson et al. (2014), Klein and Hebestreit (2015), Chen et al. (2016), Zhang et al. (2016), Yong et al. (2016), Wreczycka et al. (2017), Shafi et al. (2017), Huh et al. (2017), and Liu et al. (2020), among others.

Despite the availability of these methods, they have limitations in addressing the challenges inherent in BS-Seq data and do not provide a robust and efficient DMC/DMR identification procedure in different settings. Therefore, there is a need to develop new methods that can effectively tackle the challenges and accurately identify DMCs/DMRs.

This paper presents a novel approach for identifying DMCs in cancer epigenetic studies, using a trans-dimensional Markov chain Monte Carlo (TMCMC) with HMMs and binomial emissions in BS-Seq data. Our approach offers flexible modeling of the methylation states and transition probabilities, overcoming all known challenges inherent in BS-Seq data. Section 2 describes datasets on Colorectal cancer, whole blood cell type, and whole-genome BS-Seq (WGBS) on large offspring syndrome, which motivated our research and method development. Section 3 provides the description of our approach, including a penalized data-driven reversible jump algorithm (DDRJ), a Bayesian hierarchical model and a false discovery rate (FDR) control. Section 4 provides comprehensive data-driven simulation studies, and Section 5 presents an in-depth analysis of a Colorectal cancer dataset. We conclude with final remarks in Section 6.

## 2 Motivating Real Datasets

This research is motivated by several BS-Seq datasets described as follows.

### 2.1 Colorectal Cancer Data

We obtained reduced-representation BS-Seq (RRBS) and RNA-seq data from the NCBI Gene Expression Omnibus (GEO) with accession no. GSE95656 (Hanley et al., 2017). The data included information on 10 KRAS-muted Stage III–IV CRC samples and their adjacent normal samples which we refer to them as the **CRC** dataset, as well as 10 KRAS-mutated Aberrant Crypt Foci (ACF) samples and their normal-appearing mucosa from the distal colon of individuals without familial adenomatous polyposis or hereditary non-polyposis CRC which we refer to them as the **ACF** dataset. Only non-smokers aged between 50 and 65 were included to control for age and smoking effects on DNA methylation. Illumina HiSeq2500 was used for sequencing, with a minimum of 14 and 40 million reads per sample for RRBS and RNA-seq datasets, respectively. After removing 2,653 positions with very high read-depths (possibly due to PCR bias), the methylation data included 22,050,746 CpGs. However, the methylation information of 95.27% and 63.88% of CpGs was missing in at least one sample and in at least half of the samples, respectively. Hanley et al. (2017) removed all the positions with read-depth less than 5 from the data, which resulted in the removal of 96.59% of CpGs. This filtering may have led to an underestimation of DMRs and misleading results. In Section 5, we reanalyzed this dataset using our proposed method and compared our findings with those of Hanley et al. (2017).

### 2.2 Whole Blood Data

We used a publicly available WGBS dataset derived from peripheral blood samples and divided it into distinct blood cell subtypes: 8 CD19^+^ B-cells, 13 CD14^+^ monocytes, and 19 CD4^+^ T-cells (Cheung et al., 2017). Specifically, we extracted data related to a small genomic region located in close proximity to the *BLK* gene on human Chromosome 8, encompassing 30,440 CpGs (spanning a distance of 2 MB), which is well-known for its hypomethylation in B-cells (Kulis et al., 2015; Shokoohi et al., 2019). We used the information near the *BLK* gene to develop our simulation settings and compared several DMC/DMR methods in Examples 1 to 3 (Section 4). The dataset is referred to as **BLK**.

### 2.3 Large Offspring Syndrome Data

We obtained a subset of WGBS data from the GEO with accession no. GSE93775, which focused on large offspring syndrome (LOS), an overgrowth phenotype observed in ruminant fetuses, and was studied by Chen et al. (2017). We used the methylation data of 8 control and LOS samples on Chromosome 29 in simulation Example 4 (Section 4).

## 3 Model and Method

We propose a novel methodology for estimating the methylation profiles of subjects (e.q., patients, healthy people, cell types, etc.) using a Bayesian TMCMC approach with HMMs and binomial emissions in BS-Seq data. While the reversible jump (RJ) algorithms (Green, 1995; Richardson and Green, 1997) have been employed in HMMs (Robert et al., 2000), they often tend to under or over-estimate the number of clusters with binomial emission, as demonstrated in a simulation study detailed in Supplementary Section S2. To address this issue, we introduce a double-penalized data-driven RJ method that selects the optimal number of clusters, estimates HMM and binomial parameters, and clusters observations more accurately.

Having estimated the methylation profiles of each subject separately, we use these estimates in a Bayesian log-transformed linear (mixed) model to detect DMCs between groups of interests, such as patients versus healthy subjects or different cell types in blood, while adjusting for other covariates like age, sex, and disease history. This approach allows us to identify meaningful differences in methylation patterns associated with different groups or conditions while accounting for potential confounding factors.

### 3.1 A Hidden Markov Model for DNA Methylation Data

Let *{*(*y*_l_, m_l_), *l* = 1, …, *L}* be the vector of methylation read counts and read-depth at the *l*^*th*^ cytosine (i.e., genomic position) of a given sample. Starting with a simple model, we can assume that the sample is a realization of an HMM (*S*_l_, *Y*_l_) with *K* states (order), where *S*_l_ is the hidden path and *Y*_l_ is the emission generated based on this path. Note that *S*_l_ represents the class of propensity for the CpG site *l* to be methylated at each read instance. The path takes values in the state space *S* = 1, …, *K*, where the elements correspond to discrete methylation levels 0 ≤*θ*_1_ *< θ*_2_ *< < θ*_*K*_ ≤ 1. We assume that methylation read *Y*_l_ follows a binomial distribution conditional on the methylation state and the read-depth m_*l*_ but may vary due to the sequencing process. We ignore read-depth’s stochastic properties, however, such stochastic properties may be of interest in other settings.

Let ***S*** = (*S*_1_,, *S*_*L*_) be the hidden path for the sample of size *L*. For a fixed *K*, the model parameters are as follows:

- ***p***_0_ = (*p*_01_,, *p*_0K_), the initial probability for ***S***, where *p*_0*K*_ = Pr(*S*_1_ = *k*),
- ***P*** = *{p*_*k*_*′*_*k*_*}*, the transition matrix of ***S***, where *p*_*k*_*′*_*k*_ = Pr(*S*_l+1_ = *k*|*S*_l_ = *k*^*′*^),
- ***θ*** = (*θ*_1_ *< < θ*_*K*_), the emission parameters associated to each state *S*_l_,

where *k*^*′*^, *k* = 1,, *K*. The emission probability Pr(*Y*_*l*_ = *y*_*l*_|*θ*_*sl*_, m_*l*_) is given as *Y*_*l*_|*S*_*l*_ = *k ∼* Binomial(m_*l*_, *θ*_*k*_), *k* = 1, …, *K*. The joint distribution of ***Y*** and ***S*** is

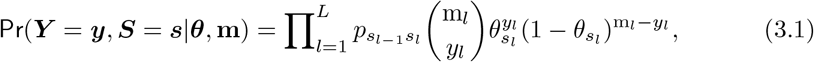

where 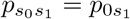. Since ***S*** is unobservable, the marginal distribution of ***Y*** is

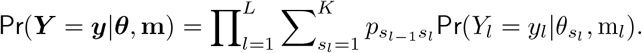

Let ***ϕ*** = (*θ*_1_, …, *θ*_*K*_, *p*_01_, …, *p*_0*K*_, *p*_11_, …, *p*_1*K*_, …, *p*_*K1*_, …, *p*_*KK*_) be the vector of parameters. The likelihood function is

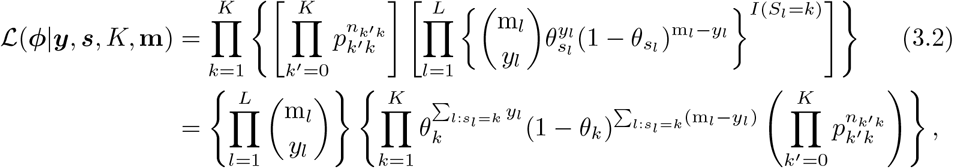

where *n*_0*k*_ = *I*(*S*_1_ = *k*) and 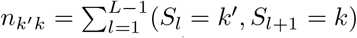 is the observed number of transitions from state *k*^*′*^ to state *k*, for all *k*^*′*^, *k* = 1, …, *K*.

### 3.2 Bayesian Trans-dimensional MCMC with HMM and Binomial

We follow a Bayesian approach similar to the ones in Zuanetti (2016), Robert et al. (2000), and Green (1995). Assume Model (3.1) where the number of states *K* is unknown and a discrete uniform distribution seems plausible for *K*. In addition, we assume *K*, ***p*** >_*k*_ and *θ*_*k*_ are independent.

Let ***ψ*** = (***ϕ***, *K*) be the vector of all parameters. The joint *a priori* distribution is

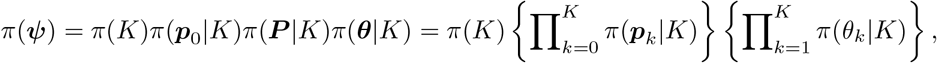

Where

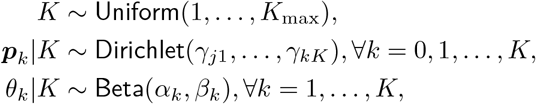

where *K*_max_ is a prespecified value, and *γ*_*kK*_, *α*_*k*_ and *β*_*k*_ are (positive) hyperparameters.

Among others, Zuanetti (2016) assumed known values for hyperparameters. In our investigation, we found out that a data-adaptive approach such as empirical Bayes increases performance and leads to faster convergence of the MCMC algorithm. To this end, we propose to either fit an inverse gamma distribution to the raw methylation levels ***β***^raw^ = **y***/***m** and estimate the shape parameter *μ* and the scale parameter *ν*, or compute the standard deviation of ***β***^raw^ and set *μ* = 1*/ν* = sd(***β***^raw^), then impose the same inverse gamma or uniform priors on the hyperparameters; that is,

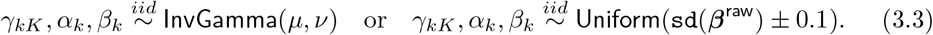

The joint *a posteriori* distribution for ***ψ*** is given as

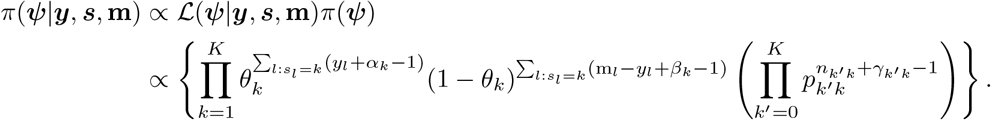

### 3.3 A Data-Driven Reversible Jump Algorithm

To select and estimate the parameters of Model (3.1), we propose a data-driven reversible jump approach in which firstly current values of parameters are updated using a Gibbs sampling procedure, and then the number of components of the HMM is updated using a split-merge move. The Gibbs sampler is basically attained by combining the likelihood function in (3.2) and obtaining the conditional *a posteriori* distributions. Accordingly, ***p***_0_ and ***p***_*k*_ are updated by generating (***γ, α, β***) using (3.3) and then

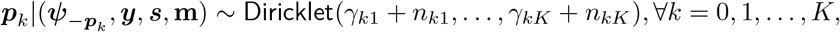

followed by predicting the path *S*_l_’s based on the conditional *a posteriori* distribution

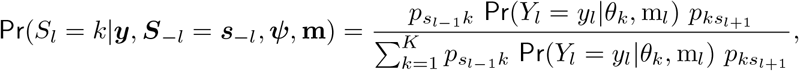

where ‘*−ω*’ means excluding *ω* in the above formulae. Evidently,

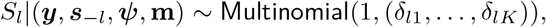

where *δ*_lk_ = Pr(*S*_*l*_ = *k*|***y, s***_*−l*_, ***ψ*, m**). The conditional *a posteriori* distribution for *θ*_*k*_ is

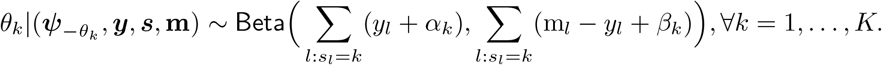

#### Split-Merge Move

To update *K*, we use trans-dimensional moves and implement a split (breaking a state in two and increasing *K* by one) and merge (collapsing two states and decreasing *K* by one) procedure. Let ***ψ*** be the current state with *K* states and ***ψ***^*\**^ be the state of the proposed movement, where *\** denotes either a split or a merge. The proposed movement is accepted based on the Metropolis-Hastings probability *ρ*(***ψ***^*\**^|***ψ***) = min(1, Λ^*\**^), where

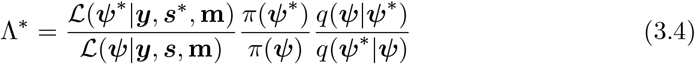

and *q*(.|.) is the proposal distribution from which we draw the proposed movement.

We apply the following algorithm on choosing a split or a merge move which is slightly different from that of Zuanetti (2016):

1. Sample with replacement two pairs of states *k*_1_ and *k*_2_ using the weights (*n*_1_, …, *n*_*k*_), where 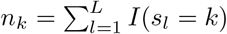.
2. If *k*_⋆_ = *k*_1_ = *k*_2_, then a split is proposed.
3. If *k*_1_ ≠ *k*_2_ and the states are neighbors, a merge of the two states is proposed. If they are not neighbors, sample without replacement until two neighboring states are selected or the number of searchers exceeds 10; then a merge is proposed.

The probability of merge is 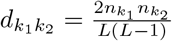, and for a split 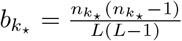, where 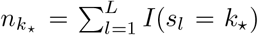 is the number of observation allocated to the new component *k*_⋆_ using simple random sampling without replacement and the arbitrary probability *ϱ*. Note that all probabilities including the probability of the new configuration ***s***^sp^ of ***S*** stay the same as in Zuanetti (2016). When *ϱ* = 0.5, we have 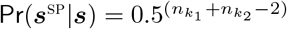.

If a split move is suggested, the parameters of the new state are estimated by drawing candidate values ***ψ***^sp^ = (*K* + 1, ***ϕ***^sp^) from the conditional *a posteriori* distribution given ***S*** = ***s***^sp^. We only need to update 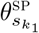 and 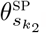. This leads to the following proposal distribution of a split:

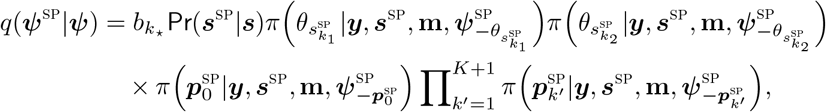

where *π*(.|.) are conditional *a posterior* distributions. We then accept the split move based on the probability min {1, Λ_sp_}. The three parts of (3.4) are given below. The likelihoods ratio for the split proposal is

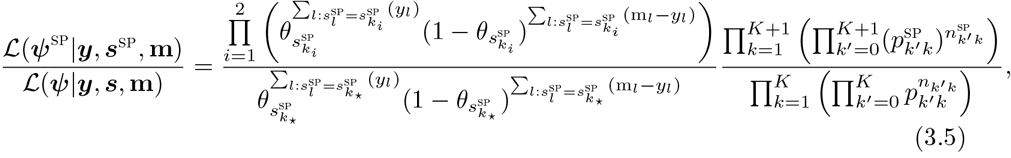

and the *a priori* distributions ratio is

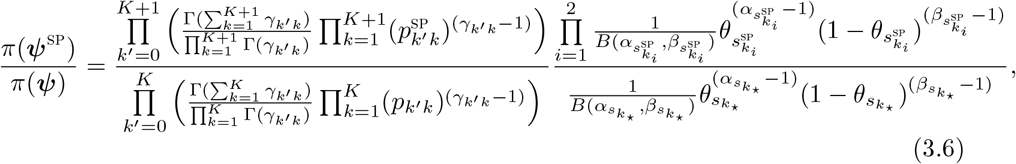

and finally, the proposal distribution ratio is

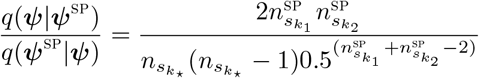

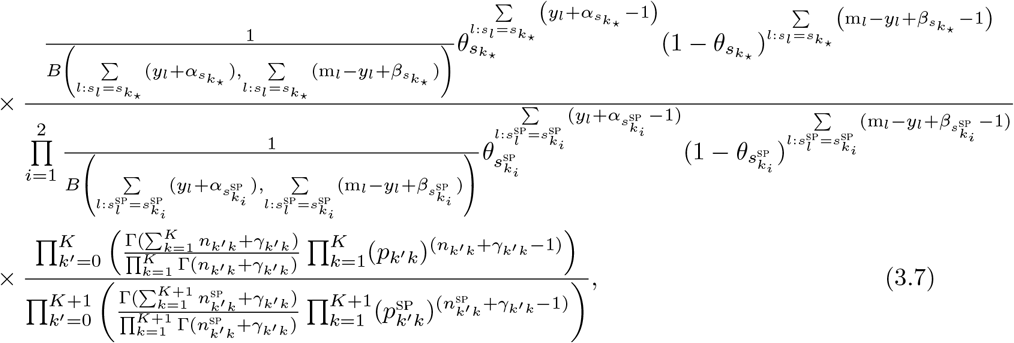

where *B*(., .) denotes the beta function. The acceptance probability for the split movement is min*{*1, Λ_sp_*}*, where Λ_sp_ is computed by multiplying (3.5), (3.6), and (3.7). Note that the acceptance probability of merging is min*{*1, Λ_mg_*}*, where Λ_mg_ = 1*/*Λ_sp_.

### 3.4 Introducing Expander-Collider Penalty

The Metropolis-Hasting probability in (3.4) and the one introduced in Zuanetti (2016) tend to over or under-estimate the number of states in Model (3.1) as demonstrated via simulations in Figures S1 to S16 in Supplementary Section S2. The underestimation mainly occurs when the number of states *K* or the sample size *L* is large. The overestimation mainly occurs when the number of states *K* is very small. To address under-estimation, we impose the penalty *κ*_sp_(*K*) = *K*^K^ on *K*; hence, we have

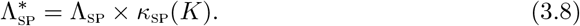

To address over-estimation, we impose the penalty 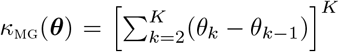 the distance between *θ*_*k*_ ‘s; hence, we have

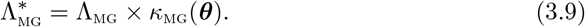

Note that *κ*_sp_(*K*) penalizes (increases) the number of states and *κ*_mg_(***θ***) ensures that two neighboring states collapse into each other when the difference between their *θ* values approaches zero. Together, these penalties constitute the *Expander-Collider* penalty function, which promotes the formation of distinct clusters and can lead to more accurate and biologically meaningful methylation profiles. Basically, the *Expander-Collider* penalty *runs with the hare and hunts with the hounds*.

The simulation results, given in Supplementary Table S3, demonstrate the superior performance of our proposed method of Expander-Collider DDRJ (ECDDRJ) over the DDRJ method in nearly all cases. We intend to avoid underestimation as it causes the merge of clusters of CpGs with different methylation patterns, thereby hindering the model’s ability to detect true differences in methylation profiles of groups of interest.

### 3.5 Bayesian Linear Mixed Model for Methylation Data

Having smoothed the methylation profile of all subjects using our proposed Expander-Collider DDRJ method described above, we collect *R* MCMC samples of parameters after a warm-up and a thining. Specifically, we store ***θ***[*s*_1:R,1_, …, *s*_1:R,L_] for each subject in the data. We then compute the average of 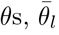, over the *R* samples and apply the logit transformation

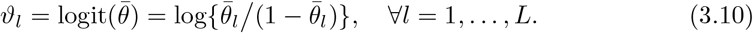

Next, we fit the following Bayesian linear (mixed) model for each CpG site *l, l* = 1, …, *L*;

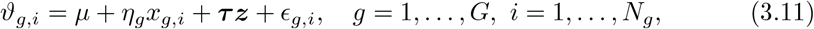

where *G* is the number of groups (e.g., *G* = 2 in the **CRC** data, *G* = 3 in the **BLK** dataset), *N*_*g*_ is the number of samples in group *g, μ* is the grand mean, *η*_*g*_ is the effect of *g*th group (*η*_1_*≡* 0), *x*_*g,i*_ = 1 if the subject belongs to group *g* and zero otherwise, ***τ*** is the vector of other (fixed and random) parameters, and ***z*** is the vector of all other covariates such as age, sex, etc. The Bayesian model is given as

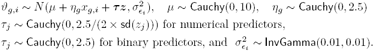

Note that Model (3.11) can be easily extended to include random effects. In our data, however, we do not have such factors.

We draw *R* MCMC samples. This can be done using the `bayesglm(.)` function in the `arm` package (Gelman et al., 2008) or the `brm(.)` function in the `brms` package (Bürkner, 2017) when random effects exist. Note that `bayesglm` gives a *p−*value for testing *H*_0_ : *η*_*g*_ = 0 which is equivalent to a *p−*value in the classical test.

#### Identifying DMCs

To detect DMCs, we may use one of the following approaches, among others:

- Apply a family-wise error control on *p*−values for testing *H*_0_ : *η*_*g*_ = 0 and estimate FDRs (Benjamini and Hochberg, 1995). FDR values that are less than a threshold level (e.g., *α* = 0.05) result in DMCs.
- Compute a 100(1−*α*)% credible interval for each *η*_*g*_. If at least one of the intervals does not include zero, the position is classified as a DMC (Shokoohi et al., 2021).
- Use global Bayesian *p*−values and Simultaneous Band Scores (Meyer et al., 2015) or Bayesian FDR (Zemplenyi et al., 2021) to detect DMCs.
- Apply the model in (3.11) on logit(*θ*_l_) on each of the MCMC samples instead of the average *𝒱* in (3.10) and obtain the proportion of times the *p*−values are significant. The position is classified as a DMC if the proportion is larger than 1 *− α*.

In our simulation studies, we found out that the first approach is the best in almost all of the settings followed by the second one. The third inflates FDR, and the last is time-consuming. Thus, the first approach is adopted here.

## 4 Simulation Study

We conducted extensive simulation studies across multiple scenarios to evaluate the effectiveness and robustness of our proposed method, which we present as Examples 1 to 4. Our objective is to demonstrate that our method outperforms other existing methods in terms of efficiency and consistency across various simulation settings.

For comparison, we selected several prominent DMC/DMR methods, including `BiSeq` (Hebestreit et al., 2013), `bumphunter` (Jaffe et al., 2012), `DMCHMM` (Shokoohi et al., 2019), `dmrseq` (Korthauer et al., 2018), `HMMDM` (Yu and Sun, 2016), `HMMDMR` (Ji, 2019), and `WFMM` (Lee and Morris, 2015). It is worth mentioning that we tested more than 20 methods (such as `HMMFisher`, `bsseq, methylKit`, `DSS`, etc.), however, their performance was not as good as the above methods. Thus, we did not report their results to save space. Our proposed method is called `DMCTHM` for simplicity.

To compare the performance of chosen methods, we compute various criteria based on the following definitions: True Positive (TP, # CpGs correctly identified as DMC); True Negative (TN, # CpGs correctly identified as non-DMC (NDMC)); False Positive (FP, # CpGs incorrectly identified as DMC); False Negative (FP, # CpGs incorrectly identified as NDMC). The sensitivity (SE) of DMRs and specificity (SP) of non-DMRs (NDMRs) are defined as 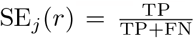 in the *j*^th^ DMR and 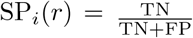 in the *i*^th^ NDMR, *j* = 1, …, *J, i* = 1, …, *J* + 1, for the *r*th simulated dataset, where *r* = 1, …, *R* = 100. The average SE_j_ of the *j*^th^ DMR and the average specificity of the *i*^th^ NDMR are calculated as 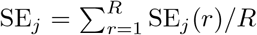 and 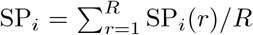, respectively.

When DMRs’ location and length are random in each simulated dataset, we only report the average overall sensitivity 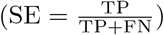, specificity 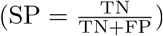, Cohen’s kappa 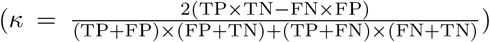, 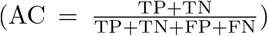 F1-score 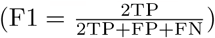, and empirical FDR (eFDR) for each method.

### 4.1 Example 1

We utilize the information near the *BLK* gene in the **BLK** dataset (Section 2). Approximately, 23.29% of the CpGs have missing information in at least one of the samples.

To generate *R* = 100 simulated datasets comprising 8 DMRs, we utilized the approach described by Shokoohi et al. (2019), which employs B-cells and monocytes. Firstly, we aggregated the read-depths and methylation counts of all B-cell samples. To impute missing information, we used nearby count information and fitted a smooth curve, G1, using `lowess` in R software with a span of 0.05, which served as the baseline profile. Then, we generated a second group curve, G2, by adding effect sizes to the chosen regions of G1 based on the information presented in Table S4 in Supplementary Section S3. To introduce additional variation, we fitted a Normal distribution to the methylation difference between G1 and all B-cell samples, obtaining estimates of *μ* = 0 and *σ* = 0.18. We then chose the read-depths of all B-cell and monocyte samples. For each dataset, we generated 8 samples from G1 and 13 samples from G2 and introduced additional variation at each site using *N* (0, 0.18) random errors, which were added to the methylation levels in G1 and G2. All values were truncated to fall within the range [0, 1]. Finally, we produced methylation counts by multiplying the generated methylation levels by read-depths and rounding to the nearest integer.

We report the results using two figures. In Figure 1, under the column named Example 1, we present the overall Sensitivity, Specificity, Accuracy, F1-score, Cohen’s kappa, and eFDR. From this figure, we observe that `DMCTHM`, `dmrseq` and `DMCHMM` give the best performance in terms of overall Sensitivity. However, `DMCTHM` outperforms both methods in terms of overall Specificity, Accuracy, Cohen’s kappa, and F1-score. Both `dmrseq` and `DMCHMM` perform poorly in terms of overall eFDR compared to `DMCTHM`.

**Figure 1:**
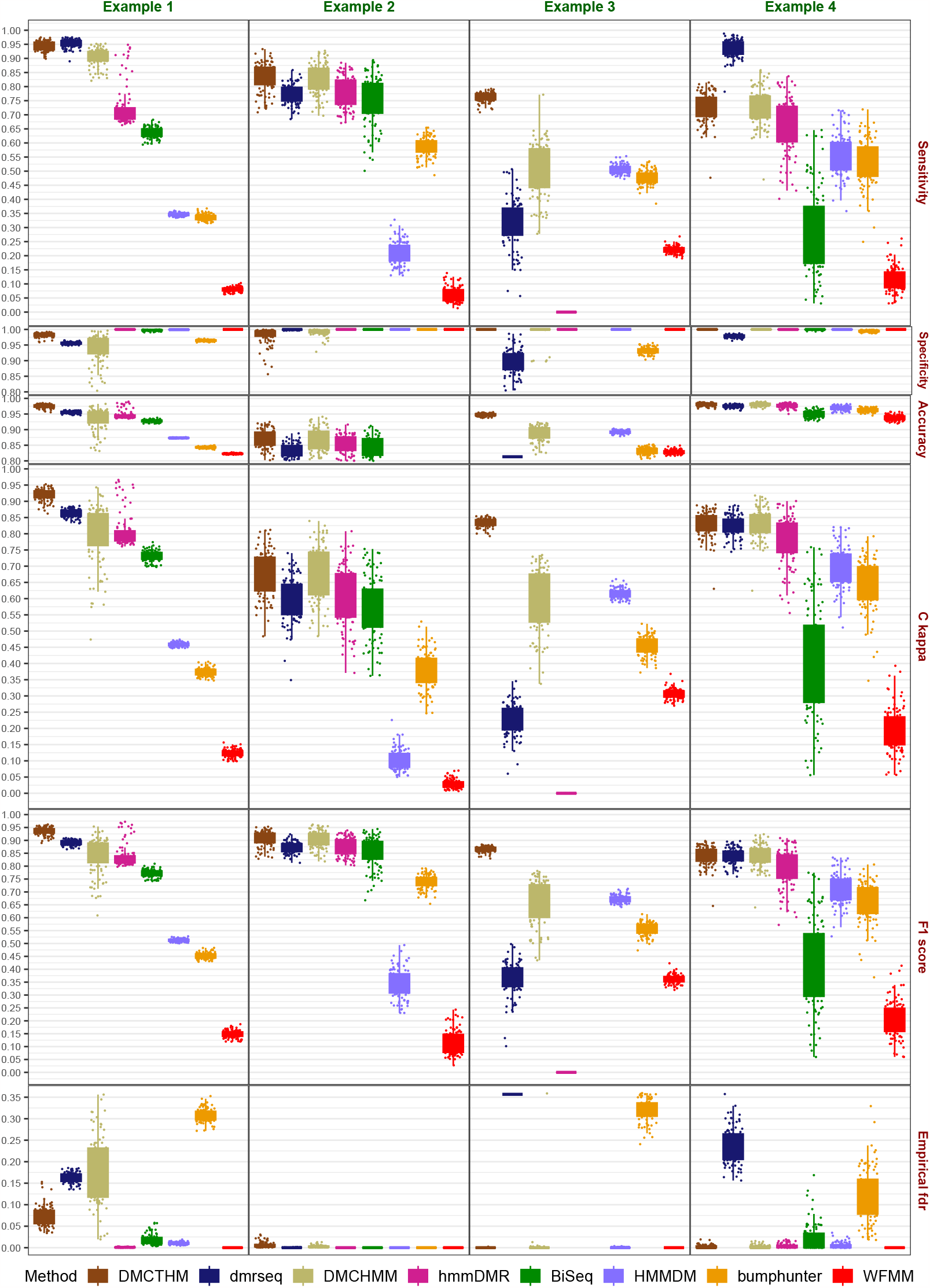
Comparison of methods for identifying DMCs in simulations using various criteria in Example 1 to 4. Except for eFDR, higher values indicate better performance.

Since the DMRs’ locations and lengths are the same in each simulated dataset, we present the Sensitivity for each DMR and ‘1 - Specificity’ for each NDMR in Figure 2. From this figure, we notice that `DMCTHM`, `dmrseq`, `DMCHMM`, and `HMMDMR` perform very well in most DMRs. However, only `DMCTHM` has the dominant Sensitivity in almost all DMRs. This is while both `dmrseq` and `DMCHMM` have higher false positive rates (1 - Specificity) compared to `DMCTHM` in all NDMRs. In general, `DMCTHM` outperforms all methods.

**Figure 2:**
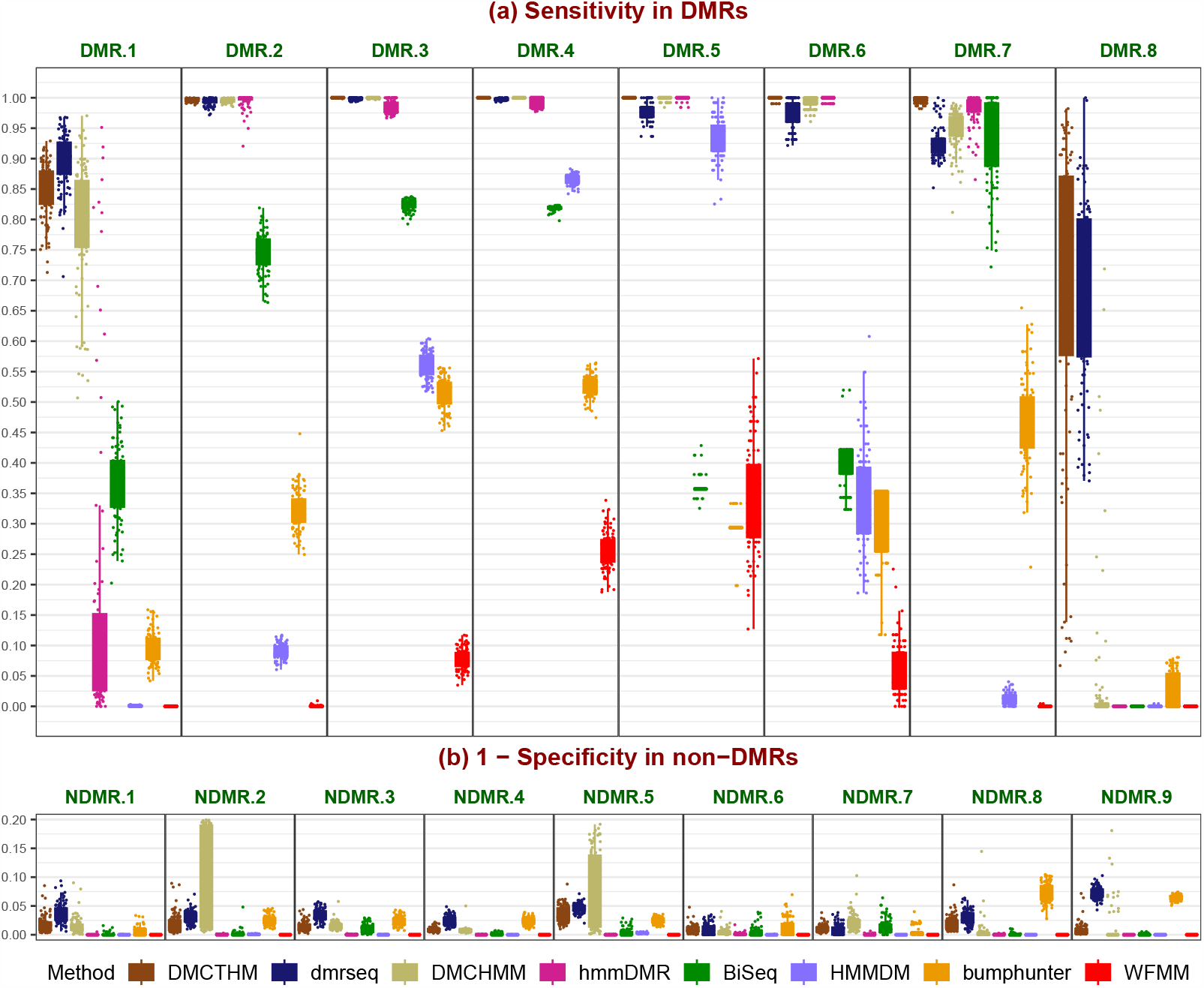
Comparison of methods for identifying DMRs in simulations using various criteria in Example 1: (a) Overall Sensitivity (True Positive) in DMRs. Higher values indicate better performance. (b) Overall ‘1 - Specificity’ (False Positive) in non-DMRs (NDMRs). Values below the nominal level of 5% are preferred.

### 4.2 Example 2

We adopted the simulation setting in Korthauer et al. (2018) by choosing 8 B-cell samples in the **BLK** dataset denoted as Group 1 and choosing the first 5,000 CpGs that do not have missing information as required by the simulation setting. A total of 50 DMRs with random lengths and locations are added to Group 1 to create Group 2. Conditional on observed coverage (read-depth), DMRs are constructed by sampling a cluster of neighboring CpGs and simulating the number of methylated reads for the samples from one population using a binomial distribution. The binomial probabilities are equal to the observed methylation proportions plus or minus an effect size that is randomly sampled from a distribution that represents small to moderate effect sizes (ranging in [0.1, 0.5]). We generate *R* = 100 datasets and computed all the criteria.

Figure 1, under the column named Example 2, depicts the results. The top five methods are `DMCTHM`, `dmrseq`, `DMCHMM`, `HMMDMR`, and `BiSeq`. Clearly, `DMCTHM` is slightly performing better. Note that this setting does not resemble a real dataset due to having no missing values. In real data between 30% to 90% of the CpGs have missing values.

### 4.3 Example 3

We follow the settings in Jaffe et al. (2012). Note that the settings are designed for microarray data by only simulating *β*-values. Thus, we modified the code to generate single nucleotide data and used the read-depths from the **BLK** dataset. We set the parameters of the simulation as follows: the number of bumps as 10, group size as 200, the number of probes in each sample as 5000, the parameter of bump maker as 0.3, the correlation structure as *AR*(1; *ϕ* = 0.21, *σ* = 0.2), and the outlier distribution as *t*_5_. To this end, we simulate 8 samples (using B-cells’ read-depths) as the baseline group and 13 samples (using monocytes’ read-depths) as Group 2 having bumps with random lengths, effects, and locations. We repeat this procedure to generate *R* = 100 datasets.

The results are presented in Figure 1 under the column named Example 3. In this setting, the `DMCTHM` method clearly dominates every other method in all criteria. Note that `BiSeq` produced errors and was not able to create any DMC/DMR in this setting.

### 4.4 Example 4

For the last example, we have adapted the simulation setting in Ji (2019). We utilized the first 5,000 CpGs of Chromosome 29 described in Section 2.3. We randomly simulated 80 DMRs with the first 40 as hypermethylated and the rest are hypomethylated and with two different lengths (500 bps and 1000 bps). For simulated DMRs, we increased and decreased the methylation rate by 0.3 for hyper- and hypo-methylated DMRs. The average methylations were truncated to stay in [0, 1]. The read-depths were kept the same in both normal and case groups. The methylation reads were generated via a Binomial distribution profile methylation levels for each sample in each group. Note that Ji (2019)’s setting only considers CpGs that have information in all samples. We revised the code to include missing values as well. We simulated *R* = 100 datasets. Figure 1, under column named Example 4, presents the results. The three top methods are `DMCTHM`, `DMCHMM`, and `HMMDMR`. Although `dmrseq` has the best overall Sensitivity, it has a very large eFDR, possibly due to aggressive smoothing.

All in all, from all the examples’ results, we can infer that `DMCTHM` is an efficient and robust method for detecting DMCs in BS-Seq data and different data settings.

## 5 Colorectal Cancer Data Analysis

We performed real data analysis on the Colorectal cancer data described in Section 2.1. Hanley et al. (2017) analyzed the data by removing all positions with missing values and read-depth lower than 5, which may have led to an underestimation of DMCs/DMRs, causing misleading results. The distribution of missing values is given in Supplementary Figure S17. We reanalyzed the data by including all CpGs using our proposed method, `DMCTHM`, and compared our results with those of the T-test in Hanley et al. (2017). In what follows, we refer to the results of Hanley et al. (2017) simply as T-test.

To proceed, we first used our proposed ECDDRJ method to predict methylation levels in both **CRC** and **ACF** datasets. We then applied the following Bayesian linear model on logit-transformed predicted methylation levels using priors described in Section 3.5 for each dataset separately:

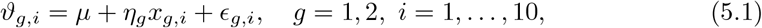

where *μ* is the grand mean, *x*_g,i_ = 0 if the sample belongs to the normal group and *x*_g,i_ = 1 otherwise, *η*_*g*_ is the effect of group *g* (by setting *η*_1_ ≡ 0), and *ϵ*_g,i_ is the random error. Finally, CpGs were classified as DMCs using the Benjamini-Hochberg FDR.

The results of our proposed method are unsurprisingly different from those of the T-test. Figure 3 and Supplementary Figures S18 and S19 depict the results. With `DMCTHM`, we uncovered 1,877,297 (8.51%) DMCs in CRC vs. normal samples in the **CRC** dataset and 108,568 (0.49%) DMCs in ACF vs. normal samples in the **ACF** dataset, which are approximately 7.96 and 8.52 times more DMCs than the T-test, respectively. Moreover, the `DMCTHM` method identified 6.17% and 4.8% of CpGs in islands and promoters as DMCs in the **CRC** data, which are 1.98 and 2.68 times higher than those of the T-test. We observe fairly similar results in the **ACF** dataset.

**Figure 3:**
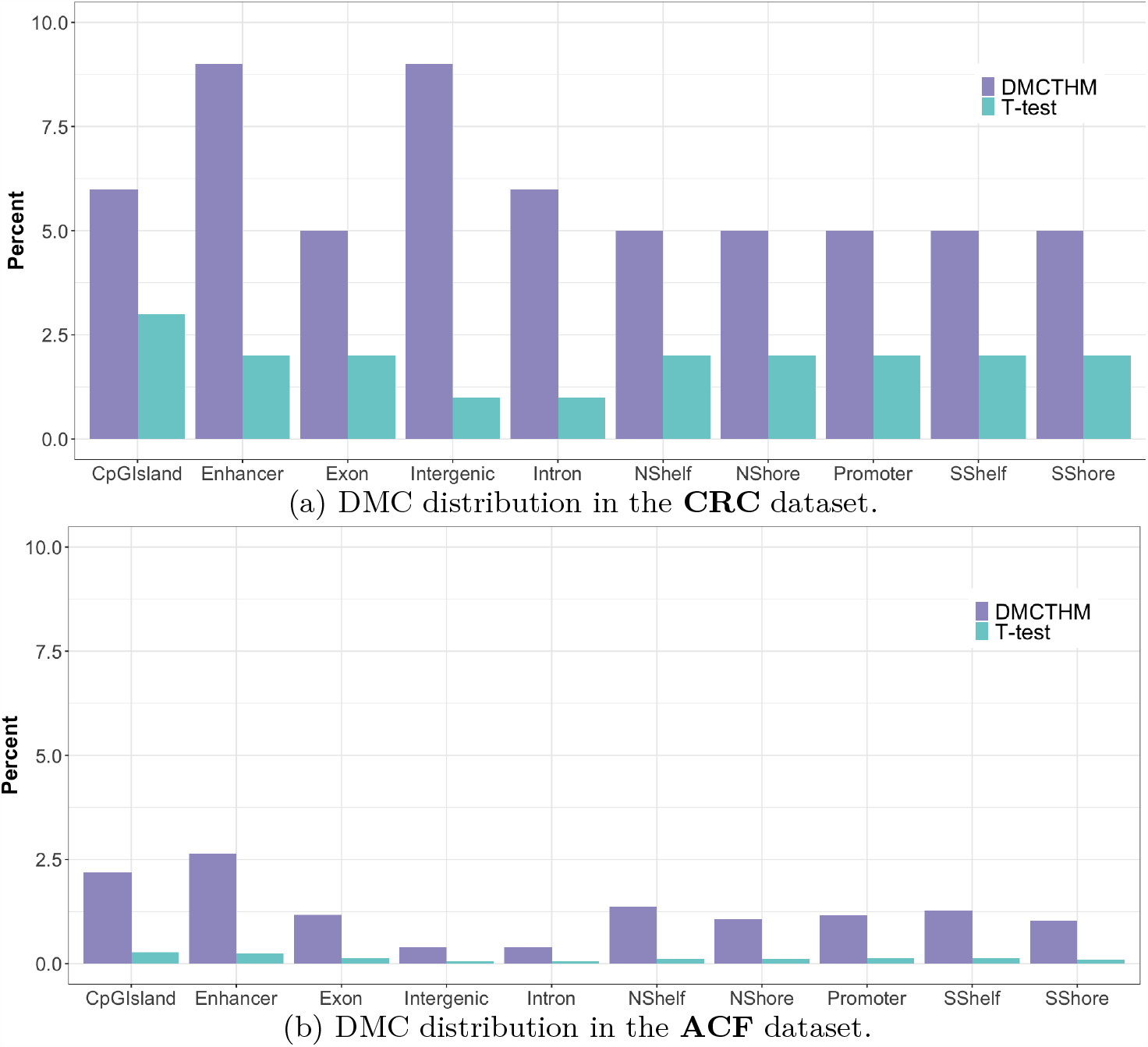
Comparison of DMC distributions in genomic regions: `DMCTHM` vs. T-test.

Figure 4 compares the different distributions of DMCs identified using `DMCTHM` in the **CRC** and **ACF** datasets. From Figure 4a, it is evident that a higher percentage of DMCs (88.9%) identified by `DMCTHM` were hypo-methylated in CRC samples compared

- DMC distribution in the **CRC** dataset.
- DMC distribution in the **ACF** dataset.

**Figure 4:**
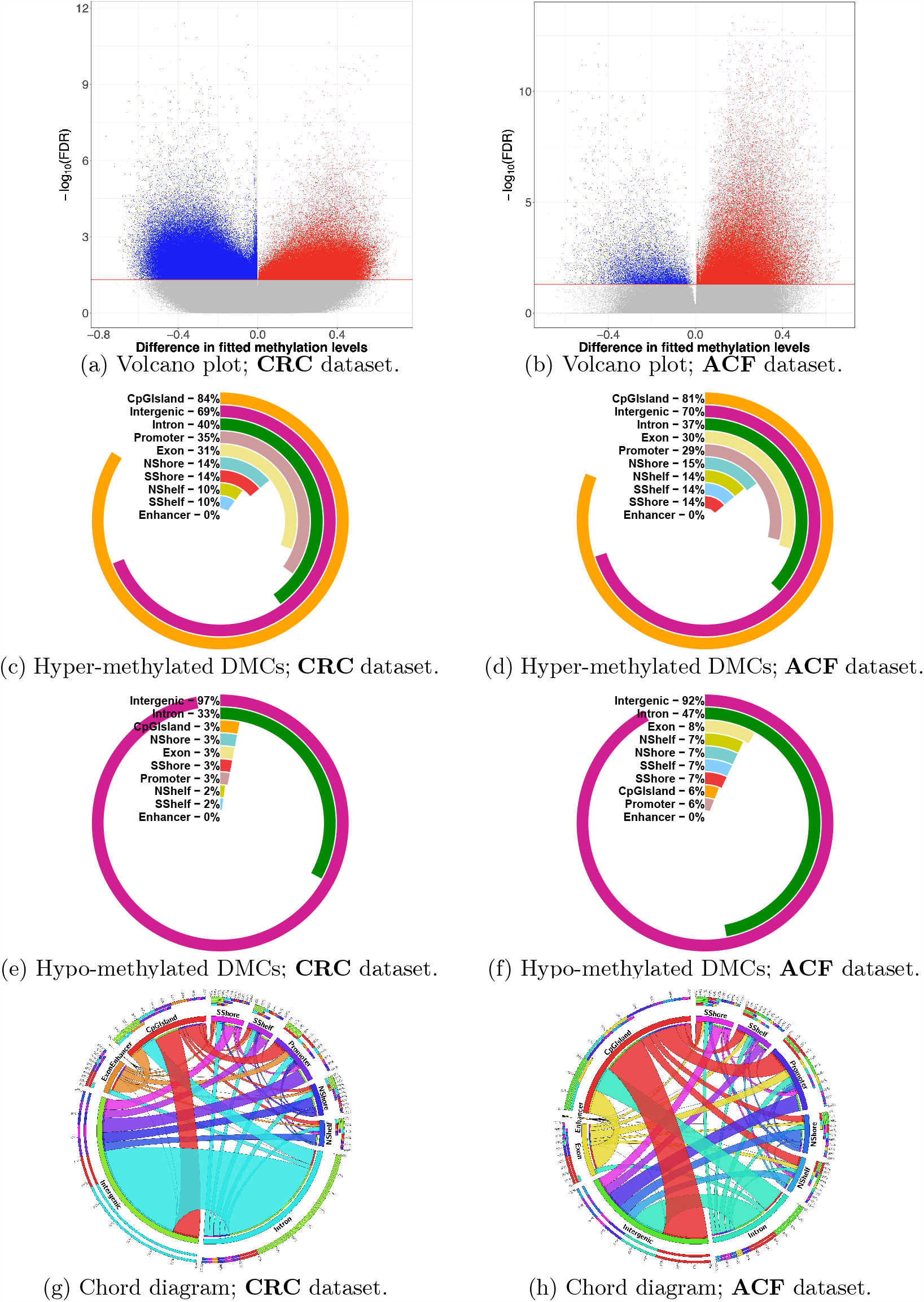
Comparison of DMC distributions obtained via `DMCTHM`.

to their adjacent normal counterparts. On the other hand, Figure 4b demonstrates that the majority of DMCs (87.9%) identified by `DMCTHM` were found to be hyper-methylated in ACF samples compared to normal crypt samples. The discrepancy in net methylation alteration between ACF and CRC indicates that a transformation in the overall methylation status might play a crucial role in the advancement from early neoplasia to invasive CRC.

From Figures 4c to 4f, the majority of hyper-methylated DMCs were located in CpG islands and intergenic regions, while hypo-methylated DMCs were primarily found in intergenic and intron regions. Furthermore, there exist almost similar patterns in DMC distribution over genomic regions in both the **CRC** and **ACF** datasets.

Figures 4g and 4h show the connections between genomic regions or locations where differentially methylated cytosines are found in the **CRC** and **ACF** datasets, respectively. From Figure 4g, we observe a very strong connection between intergenic and intron regions, and strong connections between intergenic and CpG Island, intergenic and promoter, and promoter and CpG Island, which indicate co-occurring methylation changes between these regions in the **CRC** dataset. However, the connection patterns in the **ACF** dataset are different (Figure 4h). Most notably, the connection between intergenic and intron regions is much weaker.

Table 1 shows how and when the `DMCTHM` and the T-test methods agree or disagree in the **CRC** and **ACF** datasets, respectively, with full results given in Supplementary Tables S5 and S6. The results are broken down for different genomic regions. From these tables, we observe that when the T-test identifies a CpG as a DMC, `DMCTHM` concurs with this classification 26.59% and 58.88% of the time for the **CRC** and **ACF** datasets, respectively. These proportions remain relatively consistent across all genomic locations, although they are slightly lower for the shelf region and higher for the promoter and intergenic regions compared to other regions.

**Table 1:**
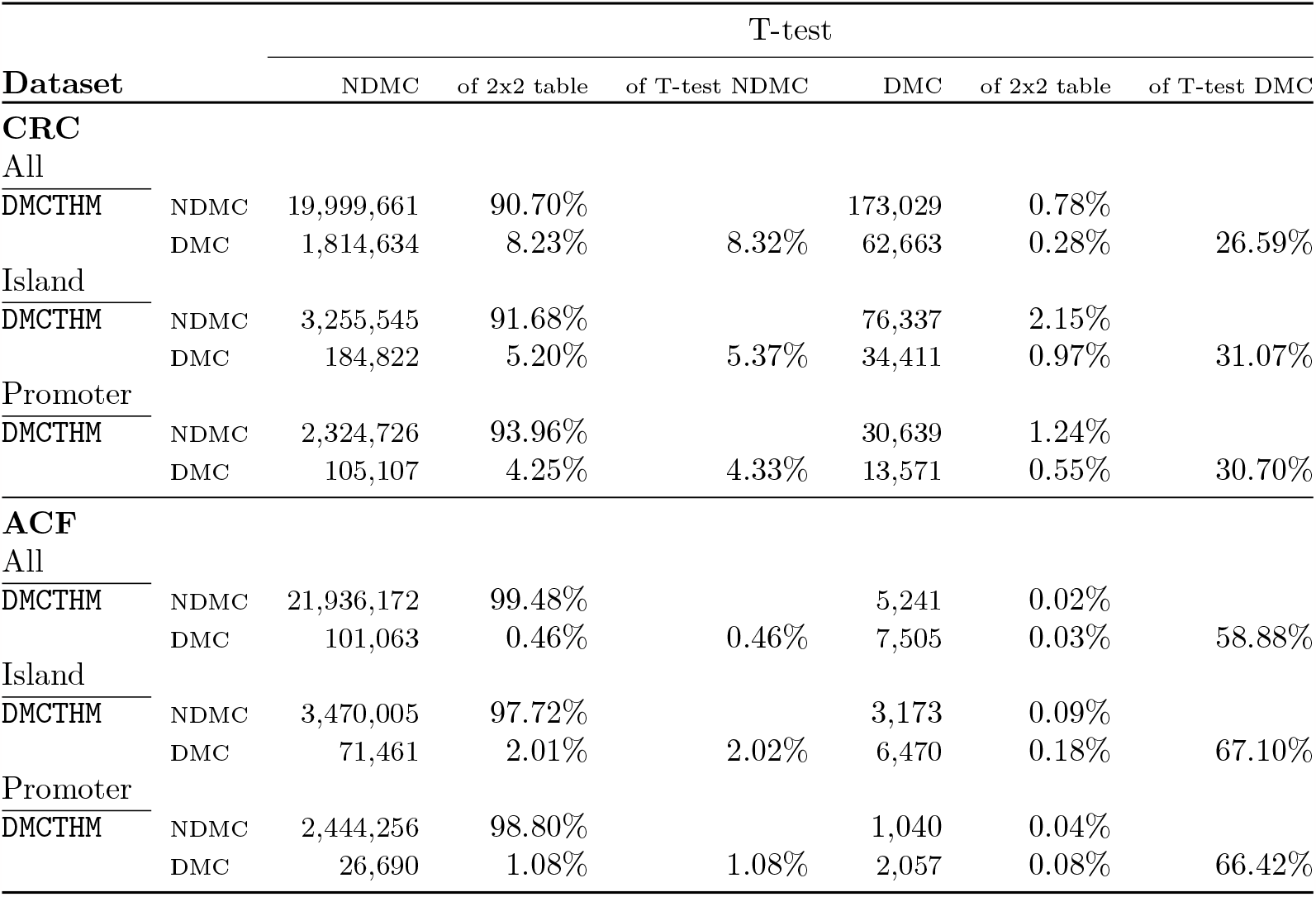
Comparison of identifying DMCs in Colorectal data: `DMCTHM` vs. T-test.

Supplementary Table S7 shows the percentage of CpGs identified as DMCs in each chromosome for both the **CRC** and **ACF** datasets. Using `DMCTHM`, we observe that the majority of DMCs were located on Chromosomes 7 (7.97%), 4 (7.62%), 2 (6.83%), 5 (6.60%), and 1 (5.21%). However, the T-test mostly located DMCs on Chromosomes 1 (7.55%), 2 (6.96%), 19 (6.78%), 7 (6.32%), and 17 (5.14%). Moreover, `DMCTHM` identified more DMCs in gene promoters in both datasets (Supplementary Table S8).

Furthermore, `DMCTHM` identified 5,699 and 932 DMGs in the **CRC** and **ACF** datasets, respectively, out of which 558 were common between the two datasets. On the other hand, the T-test identified 1,348 and 64 DMGs, out of which 50 were common among the two datasets. Moreover, `DMCTHM` and the T-test detected 924 and 59 common DMGs in the **CRC** and **ACF** datasets, respectively. Table 2 provides the full summary, including the methylation direction and the changes in direction.

**Table 2:**
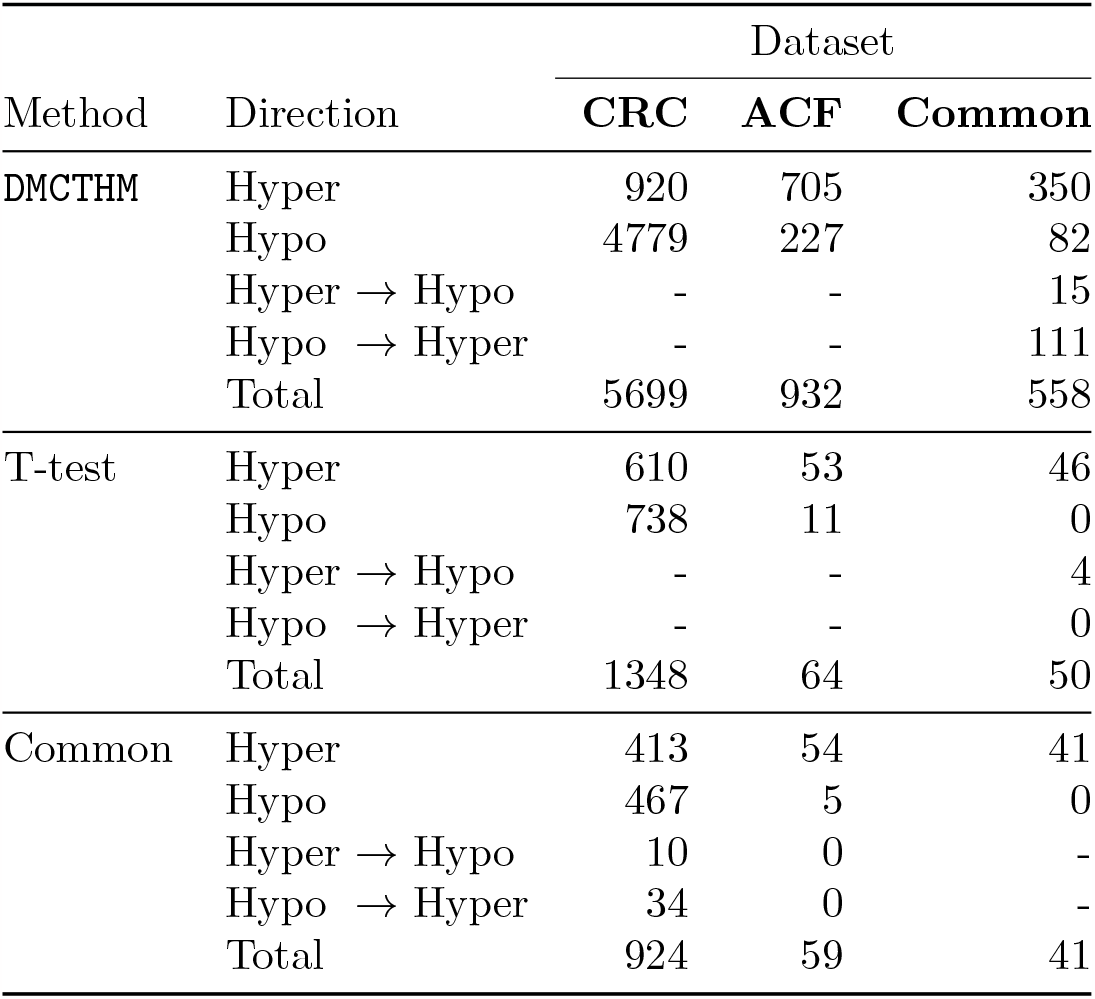
Comparison of frequency of differentially methylated genes: `DMCTHM` vs. T-test.

| Method | Direction | Dataset |  |  |
| --- | --- | --- | --- | --- |
|  |  | CRC | ACF | Common |
| DMCTHM | Hyper | 920 | 705 | 350 |
|  | Hypo | 4779 | 227 | 82 |
| | Hyper → Hypo | - | - | 15 |
| | Hypo → Hyper | - | - | 111 |
|  | Total | 5699 | 932 | 558 |
| T-test | Hyper | 610 | 53 | 46 |
|  | Hypo | 738 | 11 | 0 |
| | Hyper → Hypo | - | - | 4 |
| | Hypo → Hyper | - | - | 0 |
|  | Total | 1348 | 64 | 50 |
| Common | Hyper | 413 | 54 | 41 |
|  | Hypo | 467 | 5 | 0 |
| | Hyper → Hypo | 10 | 0 | - |
| | Hypo → Hyper | 34 | 0 | - |
|  | Total | 924 | 59 | 41 |

Additionally, Table 3 lists some of the important genes related to CRC that are identified as DMGs by `DMCTHM` and/or the T-test. The results reveal that `DMCTHM` found more CRC-related genes than the T-test, including *PMS2, CDK2A, SFRP1, SFRP5, KRAS*, and *VIM*.

**Table 3:**
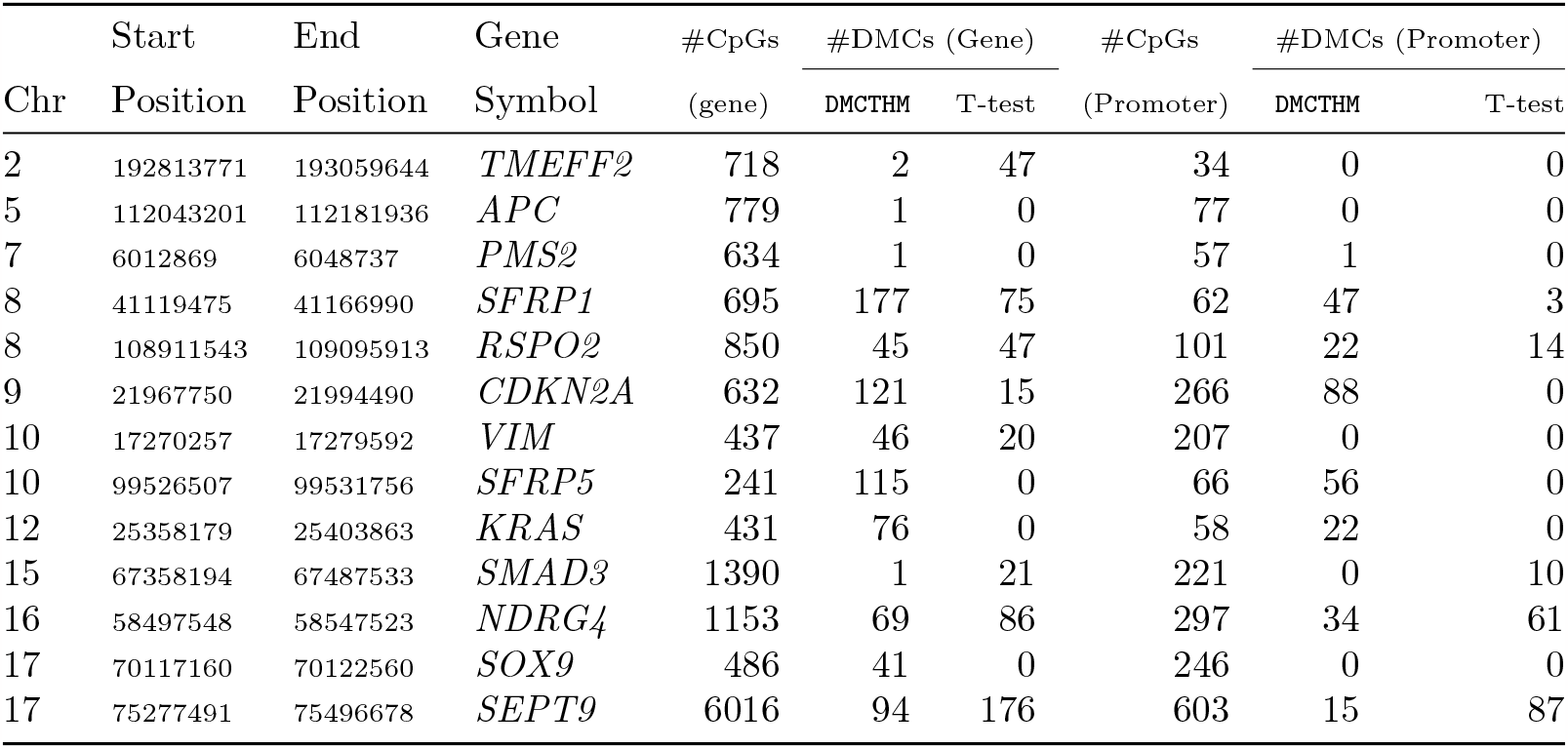
Comparison of CRC-related differentially methylated genes: `DMCTHM` vs. T-test.

| Chr | Start | End | Gene | #CpGs<br>(gene) | #DMCs (Gene) |  | #CpGs<br>(Promoter) | #DMCs (Promoter) |  |
| --- | --- | --- | --- | --- | --- | --- | --- | --- | --- |
|  | Position | Position | Symbol |  | DMCTHM | T-test |  | DMCTHM | T-test |
| 2 | 192813771 | 193059644 | *TMEFF2* | 718 | 2 | 47 | 34 | 0 | 0 |
| 5 | 112043201 | 112181936 | *APC* | 779 | 1 | 0 | 77 | 0 | 0 |
| 7 | 6012869 | 6048737 | *PMS2* | 634 | 1 | 0 | 57 | 1 | 0 |
| 8 | 41119475 | 41166990 | *SFRP1* | 695 | 177 | 75 | 62 | 47 | 3 |
| 8 | 108911543 | 109095913 | *RSPO2* | 850 | 45 | 47 | 101 | 22 | 14 |
| 9 | 21967750 | 21994490 | *CDKN2A* | 632 | 121 | 15 | 266 | 88 | 0 |
| 10 | 17270257 | 17279592 | *VIM* | 437 | 46 | 20 | 207 | 0 | 0 |
| 10 | 99526507 | 99531756 | *SFRP5* | 241 | 115 | 0 | 66 | 56 | 0 |
| 12 | 25358179 | 25403863 | *KRAS* | 431 | 76 | 0 | 58 | 22 | 0 |
| 15 | 67358194 | 67487533 | *SMAD3* | 1390 | 1 | 21 | 221 | 0 | 10 |
| 16 | 58497548 | 58547523 | *NDRG4* | 1153 | 69 | 86 | 297 | 34 | 61 |
| 17 | 70117160 | 70122560 | *SOX9* | 486 | 41 | 0 | 246 | 0 | 0 |
| 17 | 75277491 | 75496678 | *SEPT9* | 6016 | 94 | 176 | 603 | 15 | 87 |

To gain further insights into the functional implications of the identified DMCs, we conducted an over-representation analysis using the GSEA approach on the 309 DMGs identified by `DMCTHM` but not by the T-test. The results of the analysis revealed the enrichment of several cancer-related pathways, including TP53, LncRNA-mediated mechanisms of therapeutic resistance, bladder cancer, and seven other pathways (Supplementary Figure S20a). These findings suggest their potential role in the pathogenesis of Colorectal cancer. Notably, TP53 pathway showed significant enrichment (Enrichment ratio = 7.64, *p −* value = 0.027). Additionally, the hyper-methylated DMGs were found to be associated with biological processes related to neuron differentiation, cell fate commitment, and DNA-templated transcription, underscoring their relevance to the molecular mechanisms underlying CRC development and progression (Supplementary Figures S20b and S20c). Furthermore, the over-representation analysis of hypomethylated DMGs highlighted the identification of extracellular matrix components, such as the basement membrane and collagen-containing matrix, among the enriched cellular components, shedding light on their potential involvement in CRC invasion and metastasis (Li et al., 2022; Poursheikhani et al., 2020; Burtin et al., 1983).

The TP53 pathway involves the genes *CDKN2A* and *TP73. CDKN2A*, encoding the cyclin-dependent kinase inhibitor 2A, plays a crucial role in cell cycle regulation and is frequently hyper-methylated in CRC. Epigenetic alterations in this gene result in the silencing of *CDKN2A* expression, impairing its tumor suppressive function (Psofaki et al., 2010). Similarly, over-expression of *Tp73* leads to the loss of its transcriptional activity, disrupting key cellular processes such as apoptosis, cell cycle control, and DNA repair (Chen et al., 2019). Molnár et al. (2018) identified *TP73* among the top 50 hyper-methylated regions in adenomatous CRC tissues compared to normal samples.

In conclusion, our proposed `DMCTHM` method provides a more comprehensive analysis of DMCs compared to the T-test, identifying a substantially higher number of DMCs and DMGs in CRC and ACF samples. The significant differences in DMC identification between our proposed `DMCTHM` method and the T-test can be attributed to several factors. First, Hanley et al. (2017) excluded over 96% of CpGs from the analysis due to low read-depth and/or missing values. This exclusion may have led to the omission of important DMCs. Additionally, the T-test is a basic method that cannot effectively handle the common challenges encountered in DNA methylation data analysis. In contrast, the `DMCTHM` method is empowered by advanced features such as imputation of missing values, handling of low read-depth data, incorporation of multiple covariates, and consideration of autocorrelation and functional patterns in the data. These capabilities enable `DMCTHM` to efficiently identify DMCs and contribute to its superior performance compared to other methods including the T-test.

## 6 Discussion

In this paper, we have introduced a robust and efficient method, `DMCTHM`, for identifying DMCs in BS-Seq data. The method incorporates a hidden Markov model and transdimensional MCMC with binomial emission probabilities, followed by a Bayesian general linear mixed model and family-wise error control. We introduced the Expander-Collider penalty to tackle under and over-estimation in TMCMC. We have demonstrated through data-driven simulation studies and real data analysis that `DMCTHM` outperforms existing methods and consistently delivers accurate results across different simulation settings.

The method effectively handles missing values, efficiently smoothes the data, includes all data regardless of missing values or low read-depth, and accommodates additional covariates (discrete, continuous, or combinations) in the model.

Applying our proposed method to the RRBS Colorectal cancer data previously analyzed using the T-test by Hanley et al. (2017), we have uncovered a significantly greater number of DMCs, genes and their enriched pathways compared to the T-test. We have also observed genome-wide hypo-methylation in CRC samples, consistent with the advanced stages of CRC progression, supporting previous findings (Ehrlich, 2009; Hanley et al., 2017). Additionally, our results indicate that aberrant DNA methylation commonly occurs within non-transcribed intergenic regions in both ACF and CRC samples, similar to the findings in Hanley et al. (2017).

Furthermore, we performed additional analyses by identifying differentially expressed genes (DEGs) and conducting pathway and gene set enrichment analyses. Our findings revealed 1970 (365 up-regulated and 1625 down-regulated) and 47 (28 up-regulated and 19 down-regulated) DEGs in the **CRC** and **ACF** datasets, respectively (Supplementary Figure S21). In addition, distinct groups with similar gene expression profiles were observed (Supplementary Figure S22). In the **CRC** dataset, for example, all CRC samples except #54 formed a separate, cohesive cluster. The pathway analysis highlighted the involvement of several signaling pathways, including calcium signaling, cAMP signaling, and Wnt signaling, which have been extensively studied in cancer and are associated with crucial cellular processes (Supplementary Figure S23). Gene set enrichment analysis further supported the identification of DEGs related to various diseases, with biliary tract disease showing the highest enrichment score (Supplementary Figure S24).

Future improvements of the `DMCTHM` method may involve refining the Expander-Collider penalty by incorporating tuning parameters and imposing prior distributions on tuning parameters to enhance the prediction of the order in HMMs. Additionally, exploring Bayesian family-wise error control and investigating data partitioning for better identification of short DMRs can contribute to enhancing the method’s performance. We intend to create a universal pipeline by developing a package in R software using the Rcpp library and publishing it in a statistical software journal.

In conclusion, our proposed method, `DMCTHM`, provides a consistent and efficient approach for identifying DMCs. It outperforms existing methods, offers flexibility in handling various data characteristics, and yields valuable insights into the epigenetic landscape of any cancer. The method’s accuracy and efficiency make it a valuable tool for future epigenetic research and its application in diagnostic and therapeutic strategies.

## Supplementary Material

Web-based Supplementary Materials for “Uncovering Alterations in Cancer Epigenetics via Trans-Dimensional Markov Chain Monte Carlo and Hidden Markov Models” (DOI:???; .pdf). The supplementary material contains a brief literature review, several figures and tables that referred to in the main manuscript.

## Acknowledgments

The authors would like to thank Prof. Kazem Taghva, the chair of the Department of computer science, and Prof. Martin Schiller, the director of the Nevada Institute for Personalized Medicine (NIPM) at the University of Nevada-Las Vegas (UNLV), for their financial support and help. Also, the authors would like to thank Samuel Black for his help in the C programming language.

## Notes

* **Funding:** Farhad Shokoohi is supported by Start-up Grant number PG18929 and ‘In Support of Research Scholar’ Grant number PG18494, University of Nevada-Las Vegas. This research is partially supported by NIPM, the Department of Computer Science at UNLV, and the Center of Biomedical Research Excellence through COBRE Pilot Grant number P20GM121325.

### Competing Interest Statement

The authors have declared no competing interest.

## References

Akalin, A., Kormaksson, M., Li, S., Garrett-Bakelman, F., et al. (2012). “methylKit: a comprehensive R package for the analysis of genome-wide DNA methylation profiles.” Genome Biology, 13(10): R87. 2

Ashktorab, H. and Brim, H. (2014). “DNA methylation and colorectal cancer.” Current colorectal cancer reports, 10: 425–430. 2

Benjamini, Y. and Hochberg, Y. (1995). “Controlling the False Discovery Rate: A Practical and Powerful Approach to Multiple Testing.” Journal of the Royal Statistical Society: Series B (Methodological), 57(1): 289–300. 10

Bürkner, P.-C. (2017). “brms: An R Package for Bayesian Multilevel Models Using Stan.” Journal of Statistical Software, 80(1): 1–28. 10

Burtin, P., Chavanel, G., Foidart, J.-M., and Andrfi, J. (1983). “Alterations of the basement membrane and connective tissue antigens in human metastatic lymph nodes.” International Journal of Cancer, 31(6): 719–726. 19

Chen, C., Shu, L., and Zou, W. (2019). “Role of long non-coding RNA TP73-AS1 in cancer.” Bioscience Reports, 39(10). 20

Chen, D.-P., Lin, Y.-C., and Fann, C. (2016). “Methods for identifying differentially methylated regions for sequence- and array-based data.” Briefings in Functional Genomics, 15(6): 485–490. 3

Chen, Y., Kwok, C., Jiang, H., and Fan, X. (2021). “Detect differentially methylated regions using non-homogeneous hidden Markov model for bisulfite sequencing data.” Methods, 189: 34–43. 2

Chen, Z., Hagen, D. E., Ji, T., Elsik, C. G., et al. (2017). “Global misregulation of genes largely uncoupled to DNA methylome epimutations characterizes a congenital overgrowth syndrome.” Scientific Reports, 7(1): 12667. 4

Cheung, W. A., Shao, X., Morin, A., Siroux, V., et al. (2017). “Functional variation in allelic methylomes underscores a strong genetic contribution and reveals novel epigenetic alterations in the human epigenome.” Genome Biology, 18(1): 50. 4

Condon, D., Tran, P., Lien, Y.-C., Schug, J., et al. (2018). “Defiant: (DMRs: easy, fast, identification and ANnoTation) identifies differentially Methylated regions from iron-deficient rat hippocampus.” BMC Bioinformatics, 19(1): 31. 3

Dolzhenko, E. and Smith, A. (2014). “Using beta-binomial regression for high-precision differential methylation analysis in multifactor whole-genome bisulfite sequencing experiments.” BMC Bioinformatics, 15(1): 215. 3

Ehrlich, M. (2009). “DNA hypomethylation in cancer cells.” Epigenomics, 1(2): 239–259. 21

Feng, H., Conneely, K. N., and Wu, H. (2014). “A Bayesian hierarchical model to detect differentially methylated loci from single nucleotide resolution sequencing data.” Nucleic acids research, 42(8): e69–e69. 3

Gao, S., Zou, D., Mao, L., Zhou, Q., et al. (2015). “SMAP: a streamlined methylation analysis pipeline for bisulfite sequencing.” GigaScience, 4(1). 3

Gaspar, J. and Hart, R. (2017). “DMRfinder: efficiently identifying differentially methylated regions from MethylC-seq data.” BMC Bioinformatics, 18(1): 528. 3

Gelman, A., Jakulin, A., Pittau, M. G., and Su, Y.-S. (2008). “A weakly informative default prior distribution for logistic and other regression models.” The Annals of Applied Statistics, 2(4): 1360–1383. 10

Green, P. J. (1995). “Reversible Jump Markov Chain Monte Carlo Computation and Bayesian Model Determination.” Biometrika, 82(4): 711–732. 5, 6

Hanley, M. P., Hahn, M. A., Li, A. X., Wu, X., et al. (2017). “Genome-wide DNA methylation profiling reveals cancer-associated changes within early colonic neoplasia.” Oncogene, 36(35): 5035–5044. 4, 15, 20, 21

Hansen, K., Langmead, B., and Irizarry, R. (2012). “BSmooth: from whole genome bisulfite sequencing reads to differentially methylated regions.” Genome Biology, 13(10): R83. 3

Hebestreit, K., Dugas, M., and Klein, H. (2013). “Detection of significantly differentially methylated regions in targeted bisulfite sequencing data.” Bioinformatics, 29(13): 1647–1653. 3, 11

Hsiao, C.-L., Hsieh, A.-R., Lian, I.-B., Lin, Y.-C., et al. (2014). “A Novel Method for Identification and Quantification of Consistently Differentially Methylated Regions.” PLOS ONE, 9(5): e97513. 3

Huh, I., Wu, X., Park, T., and Yi, S. (2017). “Detecting differential DNA methylation from sequencing of bisulfite converted DNA of diverse species.” Briefings in Bioinformatics, 20(1): 33–46. 3

Jaffe, A., Murakami, P., Lee, H., Leek, J., et al. (2012). “Bump hunting to identify differentially methylated regions in epigenetic epidemiology studies.” International Journal of Epidemiology, 41(1): 200–209. 2, 11, 12

Ji, T. (2019). “A Bayesian hidden Markov model for detecting differentially methylated regions.” Biometrics, 75(2): 663–673. 2, 11, 14, 15

Jühling, F., Kretzmer, H., Bernhart, S., Otto, C., et al. (2016). “metilene: fast and sensitive calling of differentially methylated regions from bisulfite sequencing data.” Genome Research, 26(2): 256–262. 3

Klein, H. and Hebestreit, K. (2015). “An evaluation of methods to test predefined genomic regions for differential methylation in bisulfite sequencing data.” Briefings in Bioinformatics, 17(5): 796–807. 3

Korthauer, K., Chakraborty, S., Benjamini, Y., and Irizarry, R. (2018). “Detection and accurate false discovery rate control of differentially methylated regions from whole genome bisulfite sequencing.” Biostatistics, 20(3): 367–383. 3, 11, 12

Kulis, M., Merkel, A., Heath, S., Queiros, A., et al. (2015). “Whole-genome fingerprint of the DNA methylome during human B cell differentiation.” Nature Genetics, 47(7): 746–756. 4

Laird, P. (2010). “Principles and challenges of genome-wide DNA methylation analysis.” Nature Reviews Genetics, 11(3): 191–203. 3

Lea, A., Tung, J., and Zhou, X. (2015). “A Flexible, Efficient Binomial Mixed Model for Identifying Differential DNA Methylation in Bisulfite Sequencing Data.” PLOS Genetics, 11(11): e1005650. 3

Lee, W. and Morris, J. (2015). “Identification of differentially methylated loci using wavelet-based functional mixed models.” Bioinformatics, 32(5): 664–672. 3, 11

Li, R., Gao, X., Sun, H., Sun, L., and Hu, X. (2022). “Expression characteristics of long non-coding RNA in colon adenocarcinoma and its potential value for judging the survival and prognosis of patients: bioinformatics analysis based on The Cancer Genome Atlas database.” Journal of gastrointestinal oncology, 13(3): 1178. 19

Liu, Y., Han, Y., Zhou, L., Pan, X., et al. (2020). “A comprehensive evaluation of computational tools to identify differential methylation regions using RRBS data.” Genomics, 112(6): 4567–4576. 3

Mayo, T., Schweikert, G., and Sanguinetti, G. (2014). “M3D: a kernel-based test for spatially correlated changes in methylation profiles.” Bioinformatics, 31(6): 809–816. 3

Meyer, M. J., Coull, B. A., Versace, F., Cinciripini, P., et al. (2015). “Bayesian function-on-function regression for multilevel functional data.” Biometrics, 71(3): 563–574. 10

Molaro, A., Hodges, E., Fang, F., Song, Q., et al. (2011). “Sperm methylation profiles reveal features of epigenetic inheritance and evolution in primates.” Cell, 146(6): 1029–1041. 2

Molnár, B., Galamb, O., Péterfia, B., Wichmann, B., Csabai, I., Bodor, A., Kalmár, A., Szigeti, K. A., Barták, B. K., Nagy, Z. B., et al. (2018). “Gene promoter and exon DNA methylation changes in colon cancer development–mRNA expression and tumor mutation alterations.” BMC cancer, 18(1): 1–14. 20

Mueller, D. and Győrffy, B. (2022). “DNA methylation-based diagnostic, prognostic, and predictive biomarkers in colorectal cancer.” Biochimica et Biophysica Acta (BBA)-Reviews on Cancer, 1877(3): 1–12. 2

Park, Y., Figueroa, M., Rozek, L., and Sartor, M. (2014). “MethylSig: a whole genome DNA methylation analysis pipeline.” Bioinformatics, 30(17): 2414–2422. 3

Poursheikhani, A., Abbaszadegan, M. R., Nokhandani, N., and Kerachian, M. A. (2020). “Integration analysis of long non-coding RNA (lncRNA) role in tumorigenesis of colon adenocarcinoma.” BMC Medical Genomics, 13: 1–16. 19

Psofaki, V., Kalogera, C., Tzambouras, N., Stephanou, D., Tsianos, E., Seferiadis, K., and Kolios, G. (2010). “Promoter methylation status of hMLH1, MGMT, and CDKN2A/p16 in colorectal adenomas.” World Journal of Gastroenterology: WJG, 16(28): 3553. 20

Richardson, S. and Green, P. J. (1997). “On Bayesian Analysis of Mixtures with an Unknown Number of Components (with discussion).” Journal of the Royal Statistical Society: Series B (Statistical Methodology), 59(4): 731–792. 5

R.L., M., Baribault, C., and Ehrlich, M. (2013). “Modeling, simulation and analysis of methylation profiles from reduced representation bisulfite sequencing experiments.” Statistical Applications in Genetics and Molecular Biology, 12(6): 723–742. 2

Robert, C. P., Rydén, T., and Titterington, D. M. (2000). “Bayesian inference in hidden Markov models through the reversible jump Markov chain Monte Carlo method.” Journal of the Royal Statistical Society: Series B (Statistical Methodology), 62(1): 57–75. 5, 6

Robinson, M., Kahraman, A., Law, C., Lindsay, H., et al. (2014). “Statistical methods for detecting differentially methylated loci and regions.” Frontiers in Genetics, 5: 324. 3

Saito, Y. and Mituyama, T. (2015). “Detection of differentially methylated regions from bisulfite-seq data by hidden Markov models incorporating genome-wide methylation level distributions.” BMC Genomics, 16(Suppl 12): S3. 2

Saito, Y., Tsuji, J., and Mituyama, T. (2014). “Bisulfighter: accurate detection of methylated cytosines and differentially methylated regions.” Nucleic Acids Research, 42(6): e45. 2

Scott, C., Duryea, J., MacKay, H., Baker, M., et al. (2020). “Identification of cell type-specific methylation signals in bulk whole genome bisulfite sequencing data.” Genome Biology, 21(1): 156. 3

Shafi, A., Mitrea, C., Nguyen, T., and Draghici, S. (2017). “A survey of the approaches for identifying differential methylation using bisulfite sequencing data.” Briefings in Bioinformatics, 19(5): 737–753. 3

Shen, L., Zhu, J., Robert Li, S.-Y., and Fan, X. (2017). “Detect differentially methylated regions using non-homogeneous hidden Markov model for methylation array data.” Bioinformatics, 33(23): 3701–3708. 2

Shokoohi, F., Stephens, D., Bourque, G., Pastinen, T., et al. (2019). “A hidden Markov model for identifying differentially methylated sites in bisulfite sequencing data.” Biometrics, 75(1): 210–221. 2, 4, 11

Shokoohi, F., Stephens, D., and Greenwood, C. (2021). “Identifying Differential Methylation in Cancer Epigenetics via a Bayesian Functional Regression Model.” bioRxiv. 3, 10

Song, Q., Decato, B., Hong, E. E., Zhou, M., Fang, F., Qu, J., Garvin, T., Kessler, M., Zhou, J., and Smith, A. D. (2013). “A reference methylome database and analysis pipeline to facilitate integrative and comparative epigenomics.” PloS one, 8(12): e81148. 2

Srivastava, A., Karpievitch, Y., Eichten, S., Borevitz, J., et al. (2019). “HOME: a histogram-based machine learning approach for effective identification of differentially methylated regions.” BMC Bioinformatics, 20(1): 253. 3

Sun, S. and Yu, X. (2016). “HMM-Fisher: identifying differential methylation using a hidden Markov model and Fisher’s exact test.” Statistical Applications in Genetics and Molecular Biology, 15(1): 55–67. 2

Sung, H., Ferlay, J., Siegel, R. L., Laversanne, M., et al. (2021). “Global cancer statistics 2020: GLOBOCAN estimates of incidence and mortality worldwide for 36 cancers in 185 countries.” CA: a cancer journal for clinicians, 71(3): 209–249. 2

Wang, Z., Li, X., Jiang, Y., Shao, Q., et al. (2015). “swDMR: A Sliding Window Approach to Identify Differentially Methylated Regions Based on Whole Genome Bisulfite Sequencing.” PLOS ONE, 10(7): 1–12. 3

Wen, Y., Chen, F., Zhang, Q., Zhuang, Y., et al. (2016). “Detection of differentially methylated regions in whole genome bisulfite sequencing data using local Getis-Ord statistics.” Bioinformatics, 32(22): 3396–3404. 3

Wreczycka, K., Gosdschan, A., Yusuf, D., Grüning, B., et al. (2017). “Strategies for analyzing bisulfite sequencing data.” Journal of Biotechnology, 261: 105–115. 3

Yassi, M., S. Davodly, E., M. Shariatpanahi, A., Heidari, M., et al. (2018). “DMRFusion: A differentially methylated region detection tool based on the ranked fusion method.” Genomics, 110(6): 366–374. 3

Yong, W.-S., Hsu, F.-M., and Chen, P.-Y. (2016). “Profiling genome-wide DNA methylation.” Epigenetics & Chromatin, 9(1): 26. 3

Yu, X. and Sun, S. (2016). “HMM-DM: identifying differentially methylated regions using a hidden Markov model.” Statistical Application in Genetics and Molecular Biology, 15(1): 69–81. 2, 11

Zemplenyi, M., Meyer, M. J., Cardenas, A., Hivert, M.-F., et al. (2021). “Function-on-function regression for the identification of epigenetic regions exhibiting windows of susceptibility to environmental exposures.” The Annals of Applied Statistics, 15(3): 1366–1385. 10

Zhang, Y., Baheti, S., and Sun, Z. (2016). “Statistical method evaluation for differentially methylated CpGs in base resolution next-generation DNA sequencing data.” Briefings in Bioinformatics, 19(3): 374–386. 3

Zuanetti, D. A. (2016). “Efficient Bayesian methods for mixture models with genetic applications.” Ph.D. thesis, Department of Statistics, Federal University of São Carlos. 6, 7, 8, 9

